# Dynamics of vegetation cover and its driving factors in the Qinling-Daba Mountains on regional scale for 1986-2019

**DOI:** 10.1101/2023.03.22.533778

**Authors:** Yao Yonghui, Wang Baoguo

## Abstract

The Qinling-Daba Mountains is the transitional zone of the north and south of China. The vegetation in this area, characterized by complexity, heterogeneity and transition, is particularly sensitive to global climate change and human activities. Based on the normalized vegetation index (NDVI) data of the growing season from 1986 to 2019, which was synthesized by Landsat series satellite data on Google Earth Engine, this paper uses the methods of spatial analysis and Geo-detectors to clarify dynamics of vegetation cover and its main driving factors in the Qinling-Daba Mountains. The results show that: (1) vegetation coverage in study area shows a U-shaped NDVI distribution pattern in latitude, anti-U-shaped patterns in longitude and with altitude ascending. (2) the dynamics of vegetation coverage can be divided into two periods according to the result of MK mutation test (the breakthrough increasing period around 2005) and the trend of NDVI change: the slow increasing period with an increasing rate of 0.25%/a from 1986 to 2004 (R^2^ 0.74), and the rapid increasing period with an increasing rate of 0.30%/a from 2005 to 2019 (R^2^ 0.92). (3) Soil type, landform, vegetation type, land use type and annual average temperature are the main driving factors of vegetation dynamics in the Qinling-Daba Mountains, while annual precipitation, population density and GDP are the secondary driving factors of vegetation dynamics. The land use type and land management policies in the Qinling-Daba Mountains have a strong impact on vegetation cover change, and the climate warming in recent decades plays more important role than precipitation on the vegetation dynamics. These results are of great significance to comprehensively understand the impact of global climate warming and human activities on the natural environment of the Qinling-Daba Mountains.

## 1 Introduction

Vegetation as an important part of the terrestrial ecosystem (Piao et al., 2003), has a significant impact on global material and energy flows, carbon balance and climate stability at different temporal-spatial scales (Schimel et al., 2000; Albani et al., 2006; Liu et al., 2015). Because it is highly sensitive to environment change, thus vegetation is an important indicator for monitoring the climate change (Parmesan and Yohe, 2003). As an important composition of land surface environment, the dynamics of vegetation is mainly related to terrain and soil, climate conditions and human activities (Xu, 2018; Nemani et al., 2003). Regional vegetation change and its response on climate warming and human activities are hot topics in present research of global environmental change (Ma et al., 2012). The Qinling-Daba Mountains known as the north–south transitional zone of China and being a large-scale east-west ecological corridor (Zhang, 2019; Yu et al., 2022), is a sensitive and important region for climate change and human activities (Yao and Cui, 2022). And the vegetation of Qinling-Daba Mountains is also undergoing profound changes under the influence of the climate warming and the intervention of human activities (Yao and Cui, 2022), and it is needed to study the vegetation dynamics for regional environment research.

Normalized Difference Vegetation Index (NDVI) is recognized as one of the most effective indicators to study vegetation change, and has been widely used in the analysis of vegetation cover characteristics, dynamics and its driving factors (Fang et al., 2003; Anyamba and Tucker, 2005; Jong et al., 2011, Wang et al., 2019). There are many studies about the NDVI change in Qinling-Daba Mountains, however, most of studies focused on local or parts of the Qinling-Daba Mountains. For example, Sun et al (2009, 2010) studied the vegetation cover change in the south and north flanks of Qinling Mountains based on GIMMS and SPOT NDVI data, and found that the NDVI was decreasing, and the decreasing rate of the north flank was greater than that of the south flank; Zhang et al. (2011) studied the response of vegetation on Taibai Mountain to climate change and reported that the NDVI was highly sensitive to temperature based on NDVI of Landsat MSS, TM and ETM data. He et al (2011) studied the vegetation cover change of Micang Mountain based on SPOT ten-day NDVI data using linear regression and Hurst index and pointed out that the NDVI of Micang Mountain had an overall upward trend with a slight fluctuation from 1998 to 2009; Ren et al (2012) studied the vegetation cover change and its response to climate in Daba Mountains used SPOT NDVI data, and found that the NDVI showed a significant upward trend from 1998 to 2009, which was significantly correlated with the temperature in the month, the temperature in the previous one and two months; Cui et al (2012) analyzed the response of vegetation cover on temperature in Qinling Mountains from 2000 to 2009 based on MODIS NDVI data by linear regression, coefficient of variation and correlation analysis, and found that the vegetation cover in Qinling Mountains had an increasing trend, and the temporal stability of vegetation cover was inversely distributed with its distance from the human resident area. Although above studies figured out the NDVI changes related to climate change in the local areas, there is still lack of comprehensive understanding of vegetation cover in the Qinling-Daba Mountains on regional scale.

Recent years, more and more studies are aware of the significance of the Qinling-Daba Mountains on regional scale. Liu et al (2015) analyzed the temporal-spatial characteristics, trends and driving factors of vegetation cover in Qinling-Daba Mountains based on MODIS NDVI data from 2000 to 2014 by trend analysis, Hurst index and partial correlation analysis, and found that the vegetation cover showed a significant increase trend attributed to the change of precipitation. Chen et al (2019) analyzed the temporal-spatial variation of vegetation cover and its correlation with climate factors in Qinling-Daba Mountains using three NDVI data: GIMMS, SPOT VEG and MODIS, and found that the NDVI significantly increased from 1982 to 2017, and the change of vegetation cover was positively correlated with temperature and positively or negatively correlated with precipitation. Yao and Cui (2022) analyzed the NDVI variation trend and its spatial variation with elevation, slope, and land - use type based on annual growing season NDVI data from 1990–2019, and discussed the impact of climate warming and land-use on the vegetation cover dynamics. These studies showed that NDVI values in the Qinling-Daba Mountains significantly increased in last decades, however, the driving factors were limited to climate and human activity factors, other natural environmental factors such as soil, topography and geomorphology were rarely discussed, and the correlation between vegetation cover change and climate factors were different in above studies. That is to say, the driving factors of vegetation cover change on regional scale are needed to further study and clarify.

In order to further discover vegetation dynamics in the Qinling-Daba Mountains on regional scale and its driving factors, based on NDVI data (30 m) of the growing season from 1986 to 2019, which is synthesized by Landsat series satellite data, this paper studies the temporal-spatial dynamics of NDVI by the method of spatial analysis and its driving factors including climate, soil, geomorphology, topography, land use etc. by the Geo-detector method. The results of this paper are of great significance to comprehensively understand the impact of climate warming and human activities on the natural environment of Qinling-Daba Mountains.

## 2 Study area

The Qinling - Daba Mountains, composed of the Qinling Mountains in the north, Hanzhong Basin - Hanshui Valley in the middle, and the Daba Mountain in the south, are situated in central China (102°–114° E, 30 –36 ° N), covering a total area of about 30.60 × 104 km^2^ (Department of Geography, Shaanxi Normal University, 1989) and known as the transitional zone of China (Figure 1). It spans 1000 km in an east-west direction and 200-300 km in a south-north direction, encompassing 155 counties, 31 cities and 6 provinces of central China (Figure 1). As the north–south transitional zone in China, steep elevational gradients and the complex climate make it a biodiversity hotspot, the vegetation in this region gradually transitions from subtropical evergreen broadleaved forest to deciduous broadleaved forest from south to north, and has vertical zonality in the mountains (Liu and Lu, 1990); it is also a key habitat for rare animals, containing lots of nature reserves and national parks (Yu et al., 2022). Therefore, this area is one of the most important and concerning areas for biodiversity in China and one of the most sensitive areas for climate change and human activities (Zhang, 2019; Yao and Cui, 2022).

**Fig. 1.**
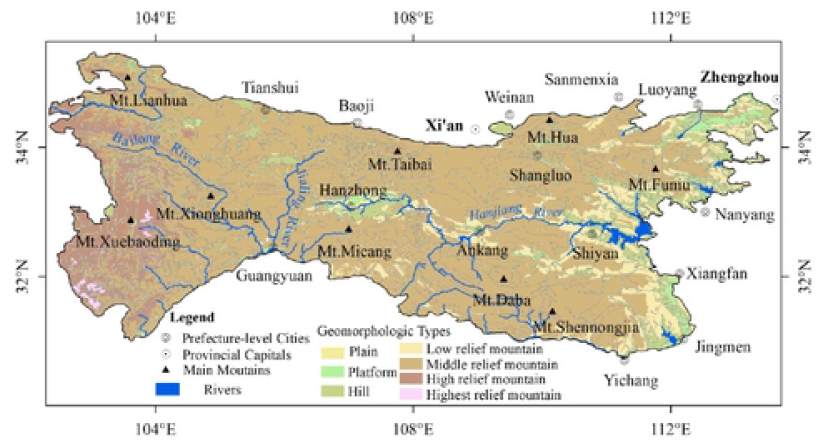
Landform map of the Qinling-Daba Mountains

## 3 Datasets and methods

### 3.1 Datasets

The NDVI dataset used in this study was the annual growing season (May to September) NDVI data (30 m resolution) of Landsat 5 / Landsat 7 / Landsat 8 from 1986 to 2019, which were synthesized using the maximum synthesis method on the Google Earth Engine (GEE) platform. Savitzky–Golay (SG) filtering was performed on the annual growing season NDVI data to further reduce the noise (Kou, 2021). Temperature and precipitation data with resolution of 500 m were downloaded from the Data Center of Resources and Environment Science, Chinese Academy of Sciences (http://www.resdc.cn) for 1980–2015, which were generated by 2400 meteorological stations using the spatial interpolation method. The ASTER GDEM (Advanced Spaceborne Thermal Emission and Reflection Radiometer Global Digital Elevation Model, downloaded from https://earthdata.nasa.gov/)with a 30 m spatial resolution were mainly used to analyze the trend of NDVI with variable altitude and slope. Land cover datasets (with 1 km resolution) in 1990, 1995, 2000, 2005, 2010 and 2015 were downloaded from the Data Center of Resources and Environment Science, Chinese Academy of Sciences (http://www.resdc.cn). Soil type data, landform data, population density and GDP data (all of them with 1 km resolution) were also downloaded from the Data Center of Resources and Environment Science, Chinese Academy of Sciences (http://www.resdc.cn). Among of them, the soil type data was generated digitally according to the “1: 1000000 Soil Map of the People’s Republic of China” compiled and published by the National Soil Census Office in 1995; the landform type data came from the Geomorphic Atlas of the People’s Republic of China (1:1000000); the population density data and GDP data were based on the statistics of population and GDP by county, using the multi-factor weight distribution method to calculate the distribution weights of land use type, night light brightness, residential area density and other related factors (Xu and Zhang, 2017). The vegetation type map data was scanned and digitized from the Vegetation Atlas of the People’s Republic of China (1:1000000) (Chinese Vegetation Map Editorial Committee, CAS, 2001), and the major categories of vegetation (coniferous forest, broad-leaved forest, mixed coniferous and broad-leaved forest, shrub, grass and other 11 categories) were extracted for the study. These data were used to analyze the influence of factors on NDVI change.

### 3.2 Methods

First, the spatial pattern and dynamics of vegetation cover was analyzed. The Sen trend method (Sen, 1968) and Mann–Kendall (MK) significant test were used to analyze the NDVI trend for 1986–2019. The Sen trend method can effectively avoids the influence of time - series data loss and data distribution form, and eliminates the interference of time - series outliers (Liu et al., 2010), and the MK significant test (Man, 1945; kendall, 1975) was conducted to test the significance of the calculated Sen trend. The MK mutation test (Karpouzos et al., 2010) were conducted to determine the mutated NDVI change period and divide the dynamic process of NDVI change. The spatial pattern of NDVI along Latitude, longitude and altitude in the Qinling-Daba Mountains were studied by profile analysis (along 33.6 ° N and 107 ° E) and statistical analysis methods.

Then the main driving factors of vegetation cover change were studied by Geo-detector. Previous studies have stated that environmental factors such as climate, terrain, landform type and soil type have a significant impact on NDVI and are the main driving factors for NDVI changes, while human activities also have important impact on NDVI changes and theirs impacts are weaker than those environmental factors (Pei et al., 2019; Yang and Han, 2019; Tao et al., 2020; Zhang et al., 2020). Therefore, 8 natural environmental factors including soil type, terrain, landform and climate, and 3 human activity factors including population, GDP and land use type (Table 1) were selected to reveal the driving factors of vegetation coverage in Qinling-Daba Mountains using the Geo-detector method based on above mentioned data every five years. As the input data of the Geo-detector requires being classified data, 7 environmental and human activity factors of elevation, slope, aspect, annual average temperature, annual precipitation, population density and GDP were classified into the classified data by the natural breakpoint method. Except the aspect, the other 6 factors were classified into 9 categories.

**Table 1.** Indexes of driving factors for NDVI change in Qinling-Daba Mountains

| Factor type | Code | Indictor | unit |
| --- | --- | --- | --- |
| Terrain | X1 | elevation | m |
|  | X2 | slope | ° |
|  | X3 | aspect | ° |
| Landform | X4 | Landform type | -- |
| Soil | X5 | Soil type | -- |
| Vegetation | X6 | Vegetation type | -- |
| Climate | X7 | Annual average type | °C |
|  | X8 | Annual precipitation | mm |
| Economy | X9 | Population density | person/km<sup>2</sup> |
|  | X10 | GDP | Ten thousand Yuan/km<sup>2</sup> |
| Land use | X11 | Land-use type |  |

The Geo-detector method was constructed on the assumption that if an independent variable has an important impact on a dependent variable, the spatial distribution of the independent variable and the dependent variable should be similar (Wang and Hu, 2011; Wang and Li, 2010; Wang and Xu, 2017). It can quantitatively expresses the spatial stratification heterogeneity of the research object by analyzing the similarities and differences between the intra-layer variance and the inter-layer variance (Hu et al., 2011; Wang et al., 2013). At present, it has been widely used to detect the driving factors in lots of studies such as land use (Hu et al., 2011), public health (Wang et al., 2013), regional economy (Ding et al., 2014), regional planning (Liu and Yang, 2012; Yang et al., 2016), meteorology and environment (Du et al., 2016) and vegetation cover change (Peng et al., 2019; Wang et al., 2019). Thus, this study uses the Geo-detector method to detect the driving factors of vegetation change.

## 4 Results and analysis

### 4.1 The temporal-spatial change of vegetation cover (NDVI)

#### (1) the spatial pattern of NDVI in the Qinling-Daba Mountains

According to the spatial distribution of the average NDVI in the Qinling-Daba Mountains from 1986 to 2019 (Fig. 2a), the vegetation coverage in the region shows a U-shaped NDVI distribution pattern in latitude and an anti-U-shaped pattern in longitude and in altitude (Fig. 2c, 2d, 2e). Mountainous areas such as the Qinling Mountains and the Daba Mountains, especially those nature reserves such as Shennongjia Nature Reserve and Taibai Mountain Nature Reserve, etc., have higher NDVI average value above 0.8 than other areas (e.g. Hanzhong basin area). Hanzhong Basin-Hanshui Valley in the middle of Qinling-Daba Mountains, the low altitude areas of Funiu Mountain, and some areas in Gansu and Sichuan Provinces have lower average NDVI value (between 0.4 and 0.6) than those mountainous areas, the NDVI value around cities along the Hanjiang River is even lower than 0.3. The NDVI in the low altitude areas in the northeast of the study area and the high altitude areas in the west is relatively low, too. The statistical analysis of NDVI average values at different altitudes shows that the NDVI first rises and then declines with the increase of altitude: the NDVI average value at altitude below 500 m is 0.6857, at altitudes between 1000 m and1500 m is 0.7884, and at altitudes between 1500 m and 3500 m are slightly declining (from 0.7743 to 0.7343); Above the altitude of 3500 m, the average NDVI value has decreased significantly, to altitude of 4000 m, the average NDVI value is declining to 0.4429 (Fig. 2e). The spatial pattern of NDVI in the Qinling-Daba Mountains shows that the landforms and elevation play important roles on the vegetation coverage.

**Fig. 2.**
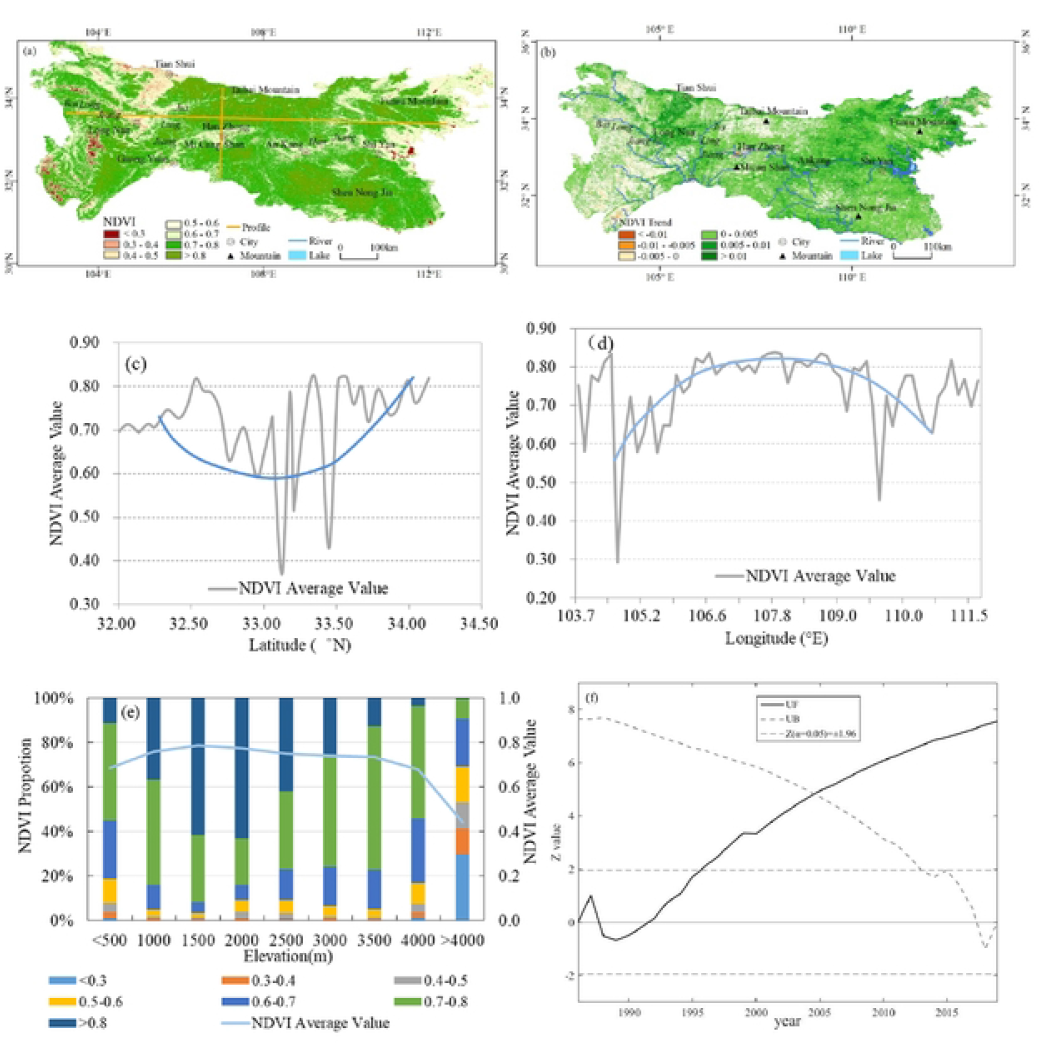
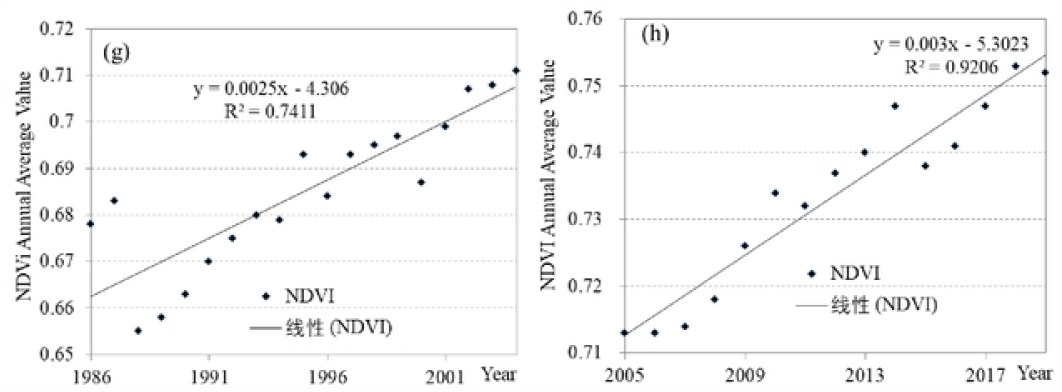
the temporal and spatial distribution of the average NDVI in the Qinling-Daba Mountains from 1986 to 2019 (a. Multi-year average NDVl from 1986 to 2019: b. Tcmporal and spatial variation of NDVI Sen trend from 1986 to 2019: c. Mul1i-year average NDVI along 107° E profile:d. Multi-year avcrage NDVI along 33.6 ° N profile; e. Molli-year averoge NDVI in altitude; f. MK mutation test of NDVI in the Qinling-Daba Mountains for 1986-2019: g. Annual avcrage NDVI trend from 1986 to 2004; h, Annual avcragc NDVI trend from 2005 to 2019)

#### (2) the temporal changes of NDVI in the Qinling-Daba Mountains

From 1986 to 2019, the vegetation cover in the Qinling-Daba Mountains showed an overall increasing trend, with an average increasing rate of 0.28%/a (R^2^ of 0.943) (Fig. 2b, 2g, 2h), indicating the continuous improvement of the vegetation cover in the study area, which was basically consistent with the previous studies (Liu et al., 2015; Chen et al., 2019; Yao et al., 2022). The distinct increasing areas of NDVI value are the places of Longnan county – Tianshui conty in Gansu Province, the west part of the Funiu Mountain and the water conservation area of the South to North Water Transfer Project, where the altitudes normally are lower than 1500m and the NDVI values are little bit lower. On the contrary, mountainous areas with altitude above 2000-3000 m especially the nature reserves with high average NDVI value have lower increasing rates and the Sen Trend values are between -0.005 and 0.005 which did not pass the significant test of 0.05 level. That is to say, the NDVI in most of mountainous areas keeps stable. The vegetation cover in the low altitude areas in the east section of the Funiu Mountain, Hanzhong Basin-Hanshui Valley, especially in the surrounding areas of cities and towns, showed a downward trend (Fig. 2b).

The result of MK mutation test on NDVI temporal series showed that NDVI had a breakthrough increasing around 2005 (Fig. 2f). Combining with the growth and development characteristics of vegetation, the dynamic process of NDVI change in the Qinling-Daba Mountains was divided into two periods: the slow increasing period with an increasing rate of 0.25%/a from 1986 to 2004 (R^2^ 0.74), and the rapid increasing period with an increasing rate of 0.30%/a from 2005 to 2019 (R^2^ 0.92).

### 4.2 Driving factors of vegetation cover change

Based on the Geo-Detector analysis for driving factors on vegetation cover change, the influence degree of each factor was ranked as follows: soil type (X5) > landform type (X4) > altitude (X1) > land use type (X11) > vegetation type (X6) > annual average temperature (X7) > slope (X2) > annual precipitation (X8) > aspect (X3) > population density (X9) > GDP (X10) (Table 2). The explanatory powers (q-value) of soil type and landform type for vegetation cover change have more than 0.2 respectively, those of altitude, land use type, vegetation type and annual average temperature are between 0.1 and 0.2 (Table 2). The above six factors are the main driving factors of vegetation cover change. The q-values of slope, annual precipitation, aspect, population density and GDP are lower than 0.1 and are secondary factors of vegetation cover change in the Qinling-Daba Mountains (Table 2).

**Table 2.** Geo-dctector analysis results of driving factors of NDVI temporal change in the Qinling-Daba Mountains

| Year | X1 | X2 | X3 | X4 | X5 | X6 | X7 | X8 | X9 | X10 | X11 |
| --- | --- | --- | --- | --- | --- | --- | --- | --- | --- | --- | --- |
| 1990 | 0.153 | 0.084 | 0.048 | 0.221 | 0.228 | 0.156 | 0.117 | 0.061 | 0.039 | — | 0.167 |
| 1995 | 0.166 | 0.100 | 0.059 | 0.242 | 0.247 | 0.172 | 0.138 | 0.045 | 0.052 | 0.028 | 0.184 |
| 2000 | 0.183 | 0.101 | 0.055 | 0.261 | 0.263 | 0.157 | 0.132 | 0.104 | 0.047 | 0.030 | 0.178 |
| 2005 | 0.150 | 0.092 | 0.058 | 0.237 | 0.259 | 0.150 | 0.131 | 0.107 | 0.036 | 0.019 | 0.185 |
| 2010 年 | 0.174 | 0.090 | 0.062 | 0.268 | 0.263 | 0.138 | 0.158 | 0.063 | 0.045 | 0.021 | 0.160 |
| 2015 | 0.207 | 0.101 | 0.036 | 0.283 | 0.268 | 0.123 | 0.175 | 0.066 | 0.047 | 0.026 | 0.146 |
| average | 0.172 | 0.095 | 0.053 | 0.252 | 0.255 | 0.149 | 0.142 | 0.074 | 0.044 | 0.025 | 0.170 |

## 5 Discussion and Conclusions

### 5.1 Discussion

#### (1) The influence of land use and climate warming on vegetation cover changes

A recent study has indicated that the “Greening Earth” is attributed to human land - use practices in China and India(Chen et al., 2019). The studies on vegetation cover change in Qinling-Daba Mountains also show that land use has a great impact on NDVI change (Sun et al., 2010; Cui et al., 2012; Yao and Cui, 2022). The results of this study also indicated that land use is also one of the main driving factors of vegetation cover change in the Qinling-Daba Mountains and its influence on NDVI change even is greater than that of climate warming (Table 2). The Qinling-Daba Mountains is not only an ecological functional area for biodiversity conservation in China, but also a water conservation area for “South to North Water Transfer Project” in China. Lots of nature reserves (over 30), national forest parks (about 37), national geological parks (11) and scenic spots (over 7) have been set up from 1960s to nowadays in the study area. The increasing of vegetation coverage is one of the achievements of those natural environment protections. Additionally, the rapid increasing of NDVI in the areas with altitudes lower than 1500 m is partly contributed to the land use policies. Chinese government issued the Grain-for-Green Policy in 1999–2000, toward which local governments formulated strict implementation measures (Chen et al., 2006; Zhang et al., 2010; Chen et al., 2019; Yao and Cui, 2022). One achievement of the Grain-for-Green was the croplands in mountainous areas with a slope steeper than 25° were required to be returned to forest (or grassland), and with slopes between 15°and 25°were conditionally returned to forest or grassland. During the field survey of the “Comprehensive Scientific Investigation of the North South Transitional Zone” project, it was also found that a large number of croplands in the mountainous areas below 1500 m were transformed into forest land again. This explains why places below 1500 m have faster NDVI increases and the breakthrough increasing period was around in 2005 (Figure 2). cropland also contributed to the NDVI increase owing to the rapidly growing hybrid cultivars, multiple croppings, irrigation, fertilizer use, pest control, improved seed quality, farm mechanization, credit availability, and crop insurance programs (Chen et al., 2019; Yao and Cui, 2022), all of which demonstrate that human land use in the Qinling - Daba Mountains has recently played a positive impact on vegetation dynamics.

#### (2) The quantification of human activity factors of vegetation dynamics

Studies about vegetation cover change in the Qinling-Daba Mountains showed that human activities play positive and negative effects on vegetation change (Deng et al., 2018; Chen et al., 2019). Land use policies, especially ecological projects such as Grain-for-Green, have a positive effect on NDVI increasing and vegetation restoration (Deng et al., 2018; Chen et al., 2019; Yao and Cui, 2022), but urbanization has also led to the NDVI decreasing and vegetation degradation in areas around the cities or resident areas (Deng et al., 2018). However, most of studies on quantitative analysis of the driving factors were focused on the climate factors and normally qualitative analysis on human activities because it was difficult to be quantified and spatialized human activity factors such as GDP and intensity of human activities. In this study, the environmental conditions, climate factors and human activity factors were selected by the Geo-detector method to comprehensively analyze the driving factors on vegetation cover change. The results also showed that land use type as one of human activities has played more important effects on vegetation cover than climate factors in the study area, and the temperature has played more important effects than precipitation (Table 2). It also states that the Geo-detector method not only can effectively and quantitatively study the driving factors of regional vegetation change, but also is helpful to the quantitative analysis of the human driving factors. Besides land use type, what factors can better reflect the impact of human activities on vegetation cover change, especially the impact of human activity intensity, remains to be further explored.

### 5.2 Conclusions

(1) Vegetation coverage with average NDVI of 0.74 in the Qinling-Daba Mountains shows a U-shaped NDVI distribution pattern in latitude, and anti-U-shaped patterns in longitude and with altitude ascending, which states that the terrain, landform, vegetation type and land use have play important roles on vegetation coverage.

(2) NDVI values in the Qinling-Daba Mountains significantly increase (with an increasing rate of 0.28%/a) and experience a dynamic change process (with a breakthrough increasing period around 2005). The process of vegetation cover change can be divided into two periods: the slow increasing period from 1986 to 2004 (with an increasing rate of 0.25%/a) and the rapid increasing period from 2005 to 2019 (with an increasing rate of 0.30%/a). The rapid increasing period were coincided with the implementation time of Grain-for-Green project and other ecological restoration projects in the early 21^st^ century, which shows that land use, especially those concerning forest conservation and expansion programs strongly contributed to the NDVI increase.

(3) Soil type, landform, vegetation type, land use type and annual average temperature are the main driving factors of vegetation dynamics in the Qinling-Daba Mountains, while annual precipitation, population density and GDP are the secondary driving factors of vegetation dynamics. The land use type and land management policies in the Qinling-Daba Mountains have a strong impact on vegetation cover change, and the climate warming in recent decades plays more important role than precipitation on the vegetation dynamics. These results are of great significance to comprehensively understand the impact of climate warming and human activities on the natural environment of the Qinling-Daba Mountains.

